# Polytypy and systematics: diversification of *Papilio* swallowtail butterflies in the biogeographically complex Indo-Australian Region

**DOI:** 10.1101/2022.03.23.485569

**Authors:** Jahnavi Joshi, Krushnamegh Kunte

## Abstract

A long-standing problem in evolutionary biology and systematics is defining patterns of diversification and speciation, which is compounded by allopatric distributions of polytypic taxa in biogeographically fragmented landscapes. In this paper we revisit this enduring systematic challenge using Mormon swallowtail butterflies (*Papilio* subgenus *Menelaides*)—an evolutionary and genetic model system. *Menelaides* is speciose and intensively sampled, with nearly 260 years of systematic study complicated by polytypy resulting from discontinuous morphological variation. This variation is structured by the mainland-island matrix of the geologically complex Indo-Australian Region, where drawing species boundaries has been difficult. We sampled variation across the biogeographic range of *Menelaides*, covering 97% of currently recognized species and nearly half of all subspecies. We generated a well-supported mito-nuclear phylogeny, on which we delineated species based on two species delimitation methods (GMYC and mPTP) and strongly supported reciprocal monophyly. These analyses showed that the true species diversity in this group may be up to 25% greater than traditional taxonomy suggests, and prompts extensive taxonomic restructuring. Biogeographic analyses showed that *Menelaides* have diversified largely in allopatry in Indo-Australian subregions by repeated dispersals across key biogeographic barriers. These results provide critical insights into the diversification process in this morphologically diverse and taxonomically complicated model group. These results will also be informative in future studies on systematics, biogeography, speciation and morphological diversification in the Indo-Australian Region—arguably the most complex geological land/seascape in the world.

## INTRODUCTION

Species are fundamental units in ecology and evolutionary biology, yet defining species boundaries has long been challenging for systematic biologists. Specifically, a key difficulty in taxonomy and systematics is defining and delineating polytypic taxa. Polytypic taxa normally arise when geographically structured, morphologically distinct but incompletely sorted lineages are identified and named as subspecies of a typically widely distributed species (Wallace 1865; Poulton 1908; Mayr 1942, 1982; de Queiroz 2020). Thus, polytypic species consist of multiple types, each defining a subspecies. Defining allopatric populations either as species or subspecies of polytypic species is taxonomically problematic. This is because reproductive isolation in such isolated populations is usually unknown, and morphological characters may vary continuously or with subtle distinction among the different allopatric populations, where delineating a species may be subjective. Large proportions of taxa of birds, fish and butterflies that have been extensively collected, studied and named are polytypic, which has created long-standing problems for taxonomists, systematists and evolutionary biologists (Mallet 2004; Andersen et al. 2014). The problem of identifying and defining polytypic species is amplified in tropical landscapes, where empirical evidence increasingly suggests the presence of cryptic and unexplored diversity across taxonomic groups (Smith et al. 2008; Joshi and Karanth 2012; Agarwal et al. 2014; Barley et al. 2015; Toussaint et al. 2015; Maddison et al. 2020). An integrative taxonomic framework, especially with multiple species delimitation methods (Sites and Marshall 2003; Pons et al. 2006; Padial et al. 2010; Fujisawa and Barraclough 2013; Zhang et al. 2013; Bouckaert et al. 2014), offers a useful tool to enumerate species diversity and to develop robust evolutionary hypotheses for polytypic taxa (Condamine et al. 2012b; Joshi and Karanth 2012; Talavera et al. 2013b, 2013a; Schwarzfeld and Sperling 2015; Toussaint et al. 2015; Matos‐Maraví et al. 2019). However, a large number of taxa even in some flagship and widely popular groups such as butterflies, beetles and odonates among the super-diverse insects, remain untouched by these approaches.

A classic group for studying geographic speciation, polytypic taxa and morphological evolution in relation to speciation is *Papilio* swallowtail butterflies, which have been extensively studied in evolutionary biology and systematics (Wallace 1865; Poulton 1908; Caterino and Sperling 1999; Caterino et al. 2001; Zakharov et al. 2004; Mallet 2004; Condamine et al. 2012b, 2012a, 2013; Allio et al. 2019; Owens et al. 2020). Sexual dimorphism, polymorphism and interbreeding of different forms of *Papilio* informed early ideas of reproductive isolation, speciation, and polytypy (Mallet 2004). Since then, the systematics of *Papilio* has been subjected to multiple, almost periodic revisions in line with the current methods and evolutionary thinking of the times, initially based purely on morphological data, but more recently with molecular data and phylogenetic methods (Munroe 1960; Hancock 1983; Caterino and Sperling 1999; Condamine et al. 2013). Recent molecular phylogenetic study on the butterfly family Papilionidae revealed that the genus *Papilio* has two distinct clades, one ranging over the Old World tropics and the other largely confined to the New World tropics (Condamine et al. 2012a, 2013). Among the seven subgenera in the Old World tropical clade of *Papilio*, subgenus *Menelaides* Hübner, [1819] is the most speciose with 56 described species and c. 200 subspecies (Häuser et al. 2005). In a recent Papilionidae phylogeny, *Menelaides* as traditionally defined was found to be non-monophyletic (Condamine et al. 2013), suggesting a need for detailed taxonomic work.

In this study, we use exhaustive taxon sampling of *Menelaides* (97% of currently recognised species and 50–60% of all known subspecies), mito-nuclear data, and detailed phylogenetic analysis to propose robust species hypotheses in an integrative taxonomic framework. We then infer biogeographic patterns and processes of diversification in this diverse Indo-Australian clade. The Indo-Australian Region is most remarkable in terms of its geological history and biological diversity among the tropical landscapes. It hosts 11 of the 33 globally recognised biodiversity hotspots consisting pieces of continental crust and oceanic islands either with Gondwanan or Eurasian origins (Hall 2012). Wallace’s seminal work in Malay Archipelago assessing the influence of geography on species distributions was important in the development of biogeographic theory (Wallace 1860, 1863). Inspired by that work, we here test the relative roles of dispersal and vicariance in influencing the current distribution of *Menelaides* within the Indo-Australian Region.

## MATERIALS AND METHODS

### Taxon Sampling and DNA Sequencing

We sampled 240 individuals representing 54 of the 56 *Papilio* (*Menelaides*) species as recently listed (Häuser et al. 2005), including ~50% of all the described subspecies (SI Table 1). The two missing *Menelaides* species were *P. lampsacus* and *P. erskinei*, which could not be sampled fresh because of their extreme rarity, and older samples did not yield DNA that could be sequenced using Sanger sequencing. All samples were either collected fresh in the field by Adam Cotton and KK or purchased from collectors by Adam Cotton, and some were donated to this study by colleagues. We extracted genomic DNA from legs and/or thoracic muscle tissue using the DNeasy Blood and Tissue Kit (Qiagen, Hilden, Germany). Polymerase chain reactions (PCR) were performed using standard primer pairs from earlier butterfly phylogenetic work (Cho et al. 1995; Brower and DeSalle 1998; Caterino and Sperling 1999; Zakharov et al. 2004; Simonsen et al. 2011). We generated approx. 4,000 bp DNA sequence data for each specimen from four mitochondrial markers (*cytochrome c oxidase I*, *cytochrome c oxidase II*, *tRNA leucine*, and ribosomal *16S* genes; total 2,623 bp) and two nuclear markers (*elongation factor I-alpha* and *wingless*; total 1,386 bp). Table S2 summarises information pertaining to each locus and primer pair. We performed PCRs with the protocol: initial 2 min denaturation at 95°C, followed by 35 cycles of 30s at 94°C, 30s at 43-60°C (depending on the primer combinations) and 1 min at 72°C, then a 4 min final extension at 72°C. We cleaned PCR products using ExoSap, and sequenced using an ABI 310 Genetic Analyzer Version 3.1 (Applied Biosystems, Foster City, USA) in the NCBS sequencing facility. Table S1 provides taxonomic and sampling details of all individuals and GenBank accession numbers of all the sequences used in this work, including the newly generated sequences (GenBank accession numbers KX557490– KX558331).

For our phylogenetic and taxonomic comparisons, we used Häuser et al. (Häuser et al. 2005) as the reference since this is the latest comprehensive catalogue of *Papilio* species. However, it should be noted that other checklists and taxonomic treatments of *Papilio* (*Menelaides*) exist (D’Abrera 1982; Beccaloni et al. 2003), which may differentially treat taxa as species or subspecies. Consequently, what we have listed as new taxonomic proposals for a more stable taxonomic arrangement in Appendix 1 based on Häuser et al. (Häuser et al. 2005) may differ with other checklists but the overall magnitude of taxonomic changes proposed and our general analyses and conclusions will remain unaffected.

### Molecular Phylogenetic Analyses

We reconstructed a molecular phylogeny of *Papilio* to ascertain the phylogenetic position of the subgenus *Menelaides* with: (a) *Menelaides* DNA sequences generated for this study (240 individuals), and (b) published sequences of 47 individuals (Zakharov et al. 2004; Condamine et al. 2013) representing all currently recognised *Papilio* subgenera: *Achillides*, *Chilasa*, *Eleppone*, *Heraclides*, *Menelaides*, *Papilio*, *Princeps*, *Pterourus*, and *Sinoprinceps.* Genus *Papilio* comprises tribe Papilionini, so we used representatives of sister tribes as outgroups: Teinopalpini (*Meandrusa payeni* and *Teinopalpus imperialis*), Troidini (*Troides helena* and *Pachliopta* spp.), and Leptocircini (*Graphium sarpedon*). We used this larger dataset with 287 individuals of all *Papilio* subgenera and seven outgroup taxa (total 294 individuals) to delineate the taxonomic position and monophyly of *Menelaides* (Fig. 2, SI Fig. 1). We subsequently used this for reconstructing the species-level phylogeny of *Menelaides* and species delimitation analyses (Fig. 3, SI Fig. 2).

**Figure 1:**
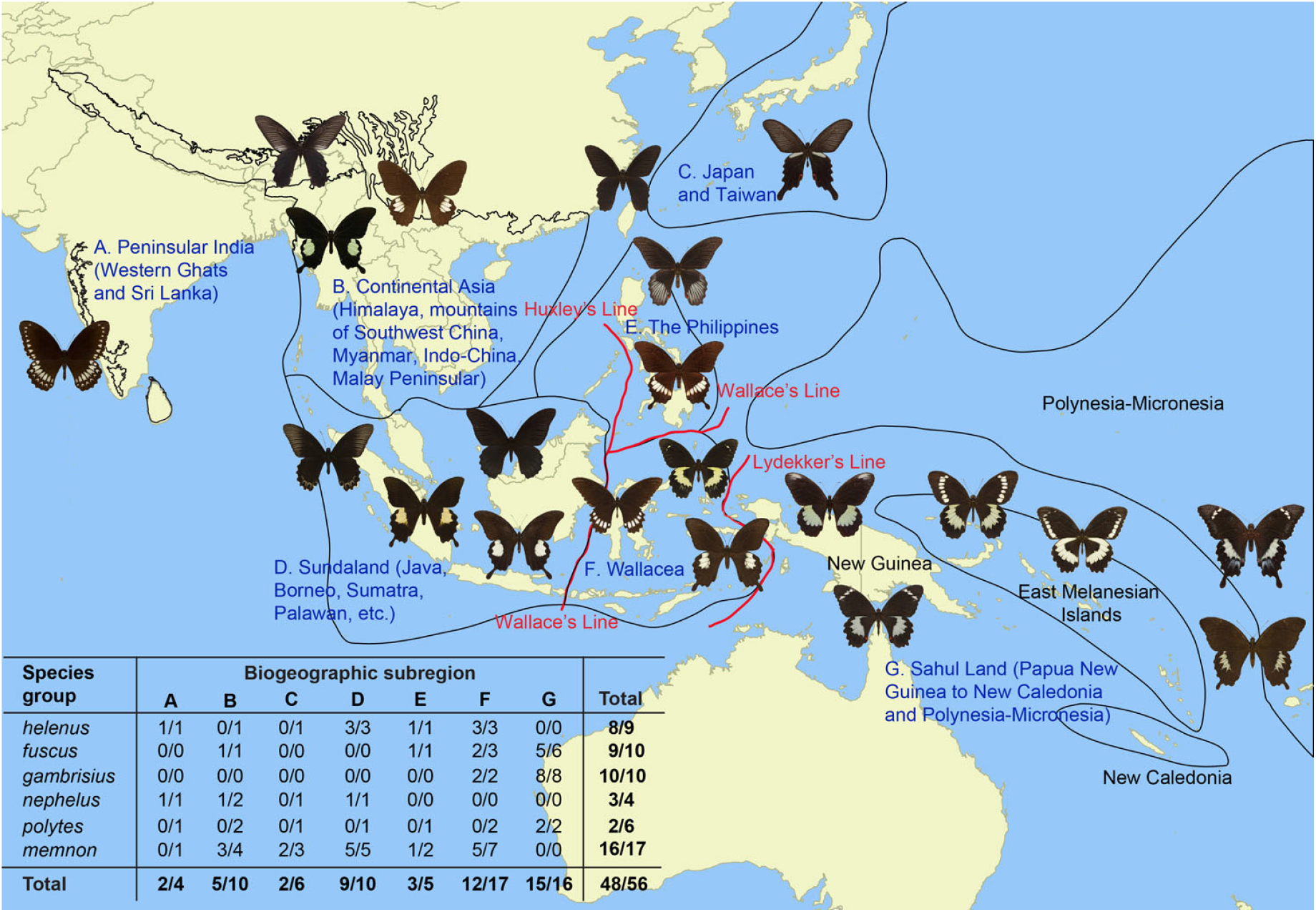
A map of the Indo-Australian Region showing globally recognised biodiversity hotspots (black outlines) and critical biogeographic barriers (red lines, and mainland-islands complexes) in the distributional range of the *Papilio* (*Menelaides*) swallowtail butterflies. We divided this landscape in seven biogeographic subregions (text in blue) as applied to distributional ranges of *Menelaides* species for biogeographic analysis. The inset table shows *Menelaides* diversity (number of endemic species/total number of species) by species groups in each biogeographic subregion, based on the total evidence presented in this paper (species delimitation analysis + strongly supported reciprocal monophyly). *Papilio erskinei* in the *gambrisius* species group (endemic to Sahul Land) and *P. lampsacus* in the *memnon* species group (endemic to Sundaland), which could not be included in this analysis, are also listed in the table.

**Figure 2:**
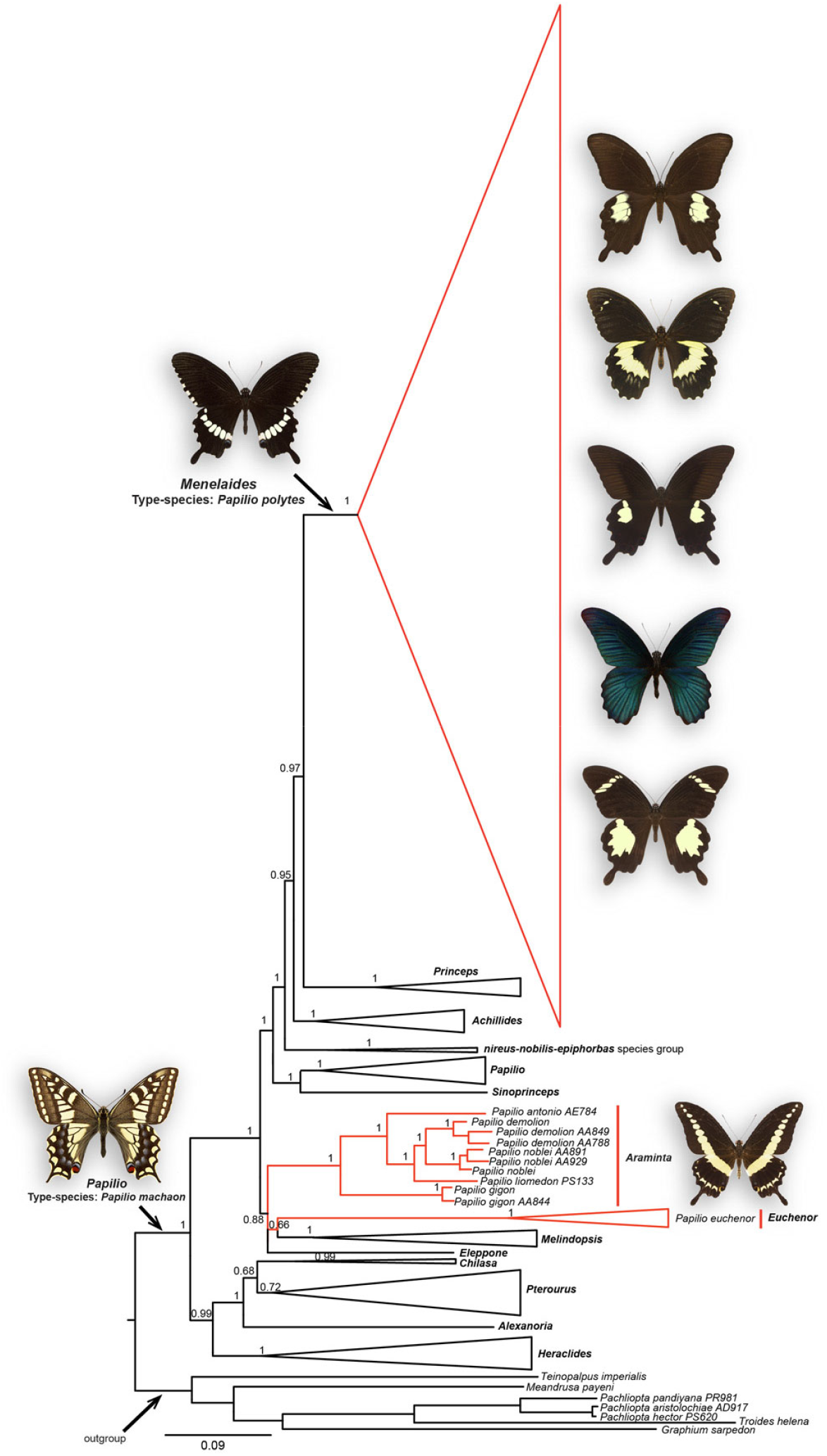
A partitioned Bayesian phylogram of *Papilio* based on four mitochondrial (*COI-tRNAleu-COII* and *16S*) and two nuclear markers (*EF1-alpha* and *Wingless*), showing the arrangement of subgenera. Bayesian posterior probability is indicated at each node. The subgenus *Menelaides* as traditionally defined was polyphyletic, which required *Menelaides* to be delineated as the most inclusive monophyletic group containing the type-species, *Papilio polytes*. Species that are taken out of *Menelaides* and reassigned to two well-supported subgenera, *Araminta* and *Euchenor*, are also marked red.

**Figure 3:**
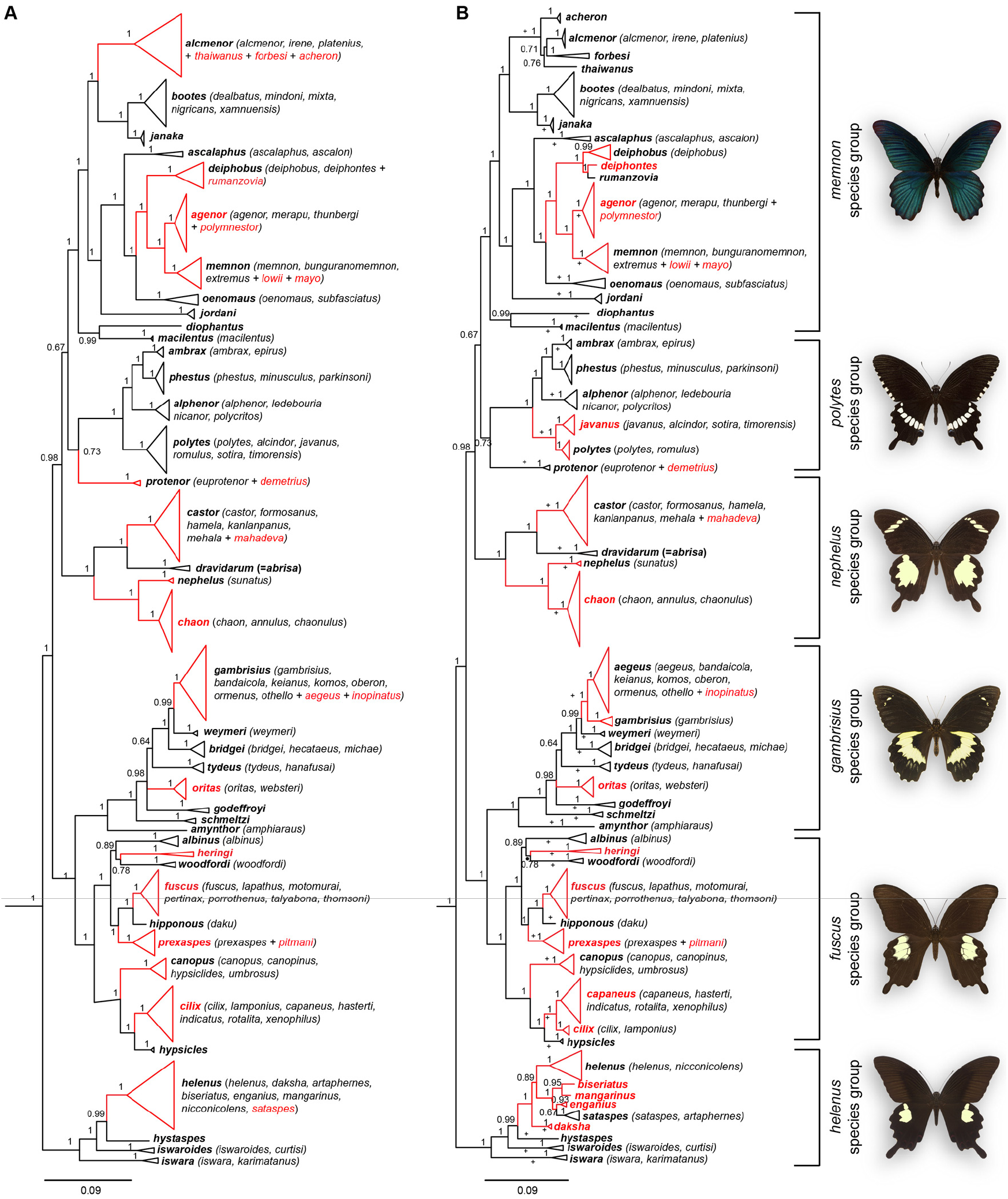
A partitioned Bayesian phylogram of the Indo-Australian *Menelaides*. **A:** Species relationships as supported by two species delimitation methods (Generalized Mixed Yule Process or GMYC, and Multi-rate Poisson Tree Process model or mPTP). **B:** Species relationships as supported by total evidence, i.e., a species delimitation method (GMYC, marked with ‘+’) and/or strongly supported reciprocal monophyly. Bayesian posterior probability is indicated at each node. A change in taxonomic status—that is, proposals for either species demoted to subspecies status or subspecies reinstated as distinct species—is proposed for the taxa that are marked in red (also see Appendix 1). Names of terminal nodes, i.e., species names as suggested by our analyses, are marked bold, whereas subspecies names in parentheses are not marked bold. *Papilio erskinei* and *P. lampsacus* are not included in this phylogeny and in Figs. 4–5 due to the lack of molecular data.

We used Bayesian and maximum likelihood approaches to reconstruct phylogenetic trees. We used PartitionFinder to choose the best partition scheme and corresponding model of sequence evolution for mtDNA and nuclear gene sequence data, using the *greedy* algorithm, where branch lenghts were linked, and models were searched for *MrBayes*, BEAST and RAxML. The Bayesian Information Criterion (BIC) was used to compare the fit of different models (Lanfear R et al. 2012). PartitionFinder suggested a total of four partitions with likelihood score of lnL −59913.96 and BIC 124976.40, specifically the mitochondrial markers 16S and *COI-COII-tRNA leu* with the GTR+I+G substitution model, the nuclear marker *EF1-alpha* with the SYM+I+G substitution model, and the nuclear marker *wingless* with the K80+I+G substitution model for MrBayes and RAxML model. For BEAST model it selected GTR+I+G substitution model for 16S and *COI-COII-tRNA leu* and HKY model for the nuclear marker *EF1-alpha*.

We performed a partitioned Bayesian analysis in MrBayes 3.2 (Ronquist et al. 2012) and set four partitions based on PartitionFinder results. We estimated base frequencies, rates for the GTR, SYM and K80 models, and the gamma distribution shape parameter, for each partition separately in MrBayes 3.2. We kept the prior distribution over the tree topologies and branch lengths at default values. The run was carried out for 50 million generations and sampled every 1,000 generations. We used split frequency below 0.01 to assess stationarity and set burn-in using MrBayes 3.2, and built a consensus tree with the remaining trees. We also checked for effective sample size (ESS) being more than 200 in Tracer v1.4.1. For maximum likelihood tree building, we used RAxML with four partitions and GTR+I, using a web-server (http://embnet.vital-it.ch/raxml-bb/) with 1,000 bootstraps. Lastly, we analysed nuclear data separately for the exhaustive dataset, using Bayesian and maximum likelihood approaches with settings as above.

### Species Delimitation Analyses

We used two widely applied species delimitation methods to define species boundaries within *Menelaides*. First, we used the Generalized Mixed Yule Coalescent (GMYC) method to identify putative species in *Menelaides* using the mitochondrial *COI-tRNA leu-COII* dataset (Pons et al. 2006). We used the R package ‘splits’ and implemented the GMYC method using both single and multi-threshold methods (Species Limits by Threshold Statistics, http://r-forge.r-project.org/projects/splits/; (Ezard et al. 2009)) to detect a threshold value for the transition from interspecific to intraspecific branching patterns and identify clusters. We performed this analysis on two datasets, one where we retained multiple individuals representing the same haplotype and another where we removed all identical haplotypes to resolve all zero-length branches. We used BEAST 2 to generate a time-calibrated ultrametric input tree (Bouckaert et al. 2014). Then we used the single threshold model with the GMYC method.

Second, we used a Bayesian implementation of the Multi-rate Poisson Tree Process model (mPTP) (Kapli et al. 2017). Higher Bayesian support value on a node indicates that all descendants from this node are more likely to be from one species. We used a single RAxML tree based on mitochondrial data for mPTP on the web-server (https://mptp.h-its.org) to run the analysis with 50,000 MCMC generations.

We also calculated pairwise genetic distance (*p*–distance) among sister-species using COI-tRNA-COII data in MEGA7 (Kumar et al. 2016).

### Divergence Time Estimation

We obtained unique haplotypes for each species defined by species delimitation analyses for mtDNA and nuDNA using Geneious 7.1.7, and used them to reconstruct a Bayesian species time tree using BEAST2 (Bouckaert et al. 2014). We used three recently estimated secondary calibrations from a well-sampled phylogenetic analysis of *Papilio*: (a) the most recent common ancestor (TMRCA) of the genus *Papilio* (mean=30.92 my, CI=25.74–38.28 my), (b) TMRCA of the Old World *Papilio* (mean=24.8 my, CI=20.69–30.68 my), and (c) TMRCA of the New World *Papilio* (mean=28.2 my, CI=23.35–35.2 my), with normal distributions (Condamine et al. 2013). In BEAST, we set the calibrations as follows: (a) *Papilio* mean=30.92 my, sigma=1, (b) Old World *Papilio* mean=24.8 my, sigma=1, and (c) New World *Papilio* mean=28.2 my, sigma=1.

We partitioned the dataset into four partitions based on PartitionFinder results, where *16S* and *COI-COII-tRNA leu* were set to GTR+I+G model and *EF1-alpha* and *wingless* to HKY substitution models. We used a relaxed molecular clock model with an uncorrelated exponential distribution and birth-death model, which has been suggested as the most appropriate model to define the speciation process (Drummond et al. 2006). For estimates of clock models, we treated the mitochondrial loci *16S* and *COI-COII-tRNA leu* as a single locus and two nuclear markers separately. We ran the program for 200 million generations and determined convergence of the chains to the stationary distribution using the program Tracer (v1.4.1) by evaluating the effective sample size. We constructed the consensus tree in TreeAnnotator (v1.4.8) and visualized it in FigTree (v1.2.2).

### Historical Biogeography Analyses

We performed historical biogeography analyses on species time-tree of *Menelaides* generated in BEAST using the R package “BioGeography with Bayesian (and likelihood) Evolutionary Analysis of RangeS (BioGeoBEARS)” (Matzke 2013). BioGeoBEARS compares alternative biogeographic models and approaches in a hypothesis-testing framework using maximum likelihood. Specifically, we performed dispersal-vicariance analysis (DIVA-like), dispersal-extinction-cladogenesis (DEC) and BayArea-like analyses in which probabilistic inference of ancestral geographical ranges and range expansion is evaluated in a maximum likelihood framework. We estimated the parameter ‘J’ to evaluate the role of founder speciation (i.e., whether dispersal leads to speciation), which is critical in the Indo-Australian Region as it is a complex mosaic of islands and continents.

We assigned each species identified as such from our species delimitation results to the following biogeographic subregions based on the distributional ranges of *Menelaides*: (A) Peninsular India, (B) Continental Asia (Himalaya, mountains of southwestern China, Myanmar, Indo-China, Malay Peninsula), (C) Japan and Taiwan, (D) Sundaland (Java, Borneo, Sumatra, Palawan and nearby islands), (E) The Philippines, (F) Wallacea (Sulawesi, the Moluccas, and the Lesser Sunda Islands), and (G) Sahul Land (Papua New Guinea, East Melanesian islands, New Caledonia, Polynesia-Micronesia) (Fig. 5). We set maximum number of subregions allowed for a species in our analysis to four based on the fact that most of the widespread species were distributed in four subregions or less. We implemented in BioGeoBEARS six models (DIVA, DIVA+J, DEC, DEC+J, BayArea, and BayArea+J) and compared likelihood scores under AIC. In this analyses, we explored the role of three processes (dispersal, vicariance and extinction), which we implemented as free parameters in a maximum likelihood framework and estimated from the data. These are maximum likelihood implementations of the original parsimony-based DIVA, likelihood-based DEC in Lagrange, and Bayesian implementation of BayArea, referred to as DIVAlike, DEClike and BayArealike (Matzke 2013). However, we will refer to them as DIVA, DEC and BayArea for simplicity.

**Figure 5:**
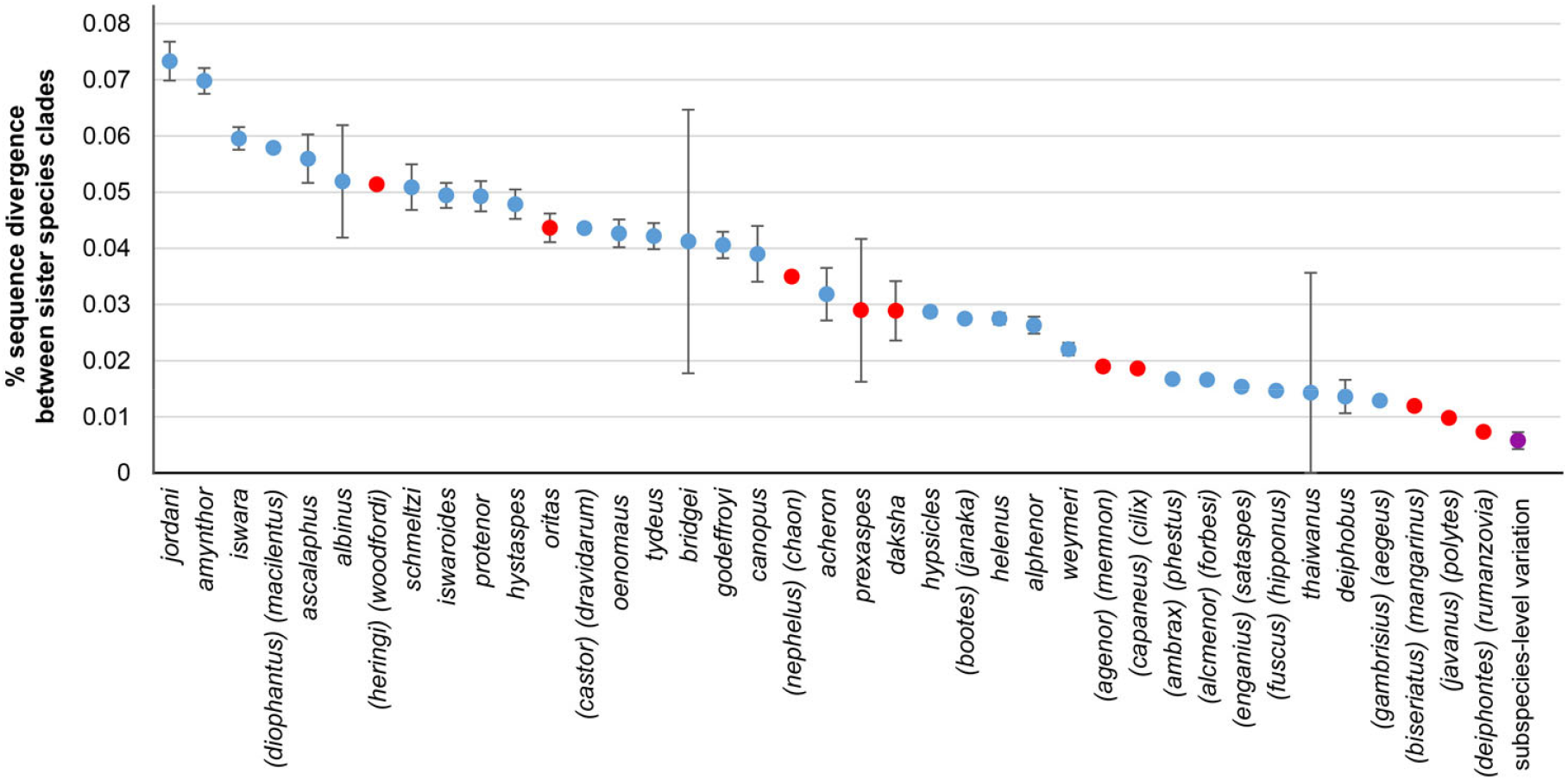
Percentage sequence divergence between sister species clades of *Menelaides* (based on Fig. 3B). If specific taxa are reciprocal sister groups of each other, then both the species names are shown (e.g., (*diophantus*, *macilentus*)). If a species has a group of species as its sister, then only the name of the species which is being compared is shown (e.g., *jordani* for (*jordani*) (*heringi*, *woodfordi*)). Blue dots show percentage divergence between sister clades that are traditionally accepted as distinct species. Red dots represent taxa elevated or reinstated in this study from subspecies to species level based on Bayesian and Maximum Likelihood phylogenetic and species delimitation analyses (see Fig. 3 and Appendix 1). Error bars are computed from multiple comparisons in cases where a single species is sister to a group of species, e.g., *P. jordani*. Shown at the right end is subspecies-level molecular divergence averaged across all the species for which multiple subspecies were sequenced.

## RESULTS

### Circumscription of *Papilio* subgenera

The subgenus *Menelaides* was polyphyletic in the partitioned Bayesian and maximum likelihood phylogenetic analyses based on 294 individuals of all the *Papilio* species, subgenera and outgroups with approx. 4kb of mitochondrial and nuclear gene sequences. Therefore, we first circumscribed *Menelaides* as the well-supported, most inclusive monophyletic clade containing the type-species, *Papilio polytes* (Fig. 2, SI Fig. 1). Six species (*antonio*, *demolion*, *noblei*, *liomedon*, *gigon* and *euchenor*) traditionally considered to be part of *Menelaides* formed two well-supported clades well outside *Menelaides* (Fig. 2, SI Fig. 1), which we propose to include in two available but rarely used subgenus names: (1) *Araminta* Moore, 1886; type-species: *Papilio demolion* Cramer, 1776 (containing *antonio*, *demolion*, *noblei*, *liomedon*, and *gigon*), and (2) the monobasic *Euchenor* Igarashi, 1979; type-species: *Papilio euchenor* Guérin-Méneville, [1830].

*Menelaides*—as delineated—was sister to the well-supported African subgenus *Princeps* Hübner, [1807]. However, *Princeps* as traditionally classified itself was polyphyletic, consisting of three distinct monophyletic clades, as reported earlier (Condamine et al. 2013). We therefore identified the sister of *Menelaides* as the true *Princeps*, consisting of the type-species *Papilio demodocus* Esper, 1799 (Fig. 2, SI Fig 1). We designated the second monophyletic group that was previously under *Princeps* as the “*nireus-nobilis-epiphorbas* species group”, rather than assign it to a subgenus name. This is because the reduced dataset of nuclear markers did not support this relationship (SI Fig. 1), indicating that better taxon sampling may be required before this species group may be formally assigned to a subgenus. We propose the available but rarely used subgenus name *Melindopsis* Aurivillius, 1899 (type-species: *Papilio rex*), for the third monophyletic group that was previously under *Princeps* (Fig. 2). Overall, these relationships were congruent in two different analyses that were based on the comprehensive mitochondrial and nuclear dataset (Fig. 2) or a trimmed dataset containing only nuclear markers (SI Fig 1). This congruence suggested that the monophyletic groups we delineated as subgenera *Menelaides*, *Princeps*, *Melindopsis*, and *nireus-nobilis-epiphorbas* species group—and their relationships with other *Papilio* subgenera—were robust against the choice of markers. This detailed molecular phylogenetic analysis and formal application of subgenus names now help stabilise the subgenera of *Papilio* and species within them.

### Phylogenetic Relationships and Systematics of *Menelaides*

We generated a robust phylogenetic hypothesis for species groups, species and subspecies within *Menelaides* by analysing our dataset in two different ways. First, we analysed the complete nuclear and mitochondrial dataset of all 294 specimens using partitioned Bayesian and maximum likelihood phylogenetic methods, where delineating species was not always straightforward (SI Fig. 2). Second, we adopted an integrative taxonomic framework in which we first analysed the complete mito-nuclear dataset of all the specimens with two species delimitation methods (Fig. 3A), and then compared geographic distributions of each strongly supported monophyletic clade in relation to known biogeographic barriers (Figs. 3b, 4). This approach resulted in a well-supported phylogeny of *Menelaides* that defined all the ‘species’ clusters from SI Fig. 2, and which also agreed closely with the expectations of taxonomic divergence based on major biogeographic barriers (Fig. 3–4). We propose this arrangement as new hypotheses for species within *Menelaides*, which supports some classical groupings but at the same time provides novel insights into species diversity in this subgenus (Fig. 3, Appendix 1), as summarised below.

**Figure 4:**
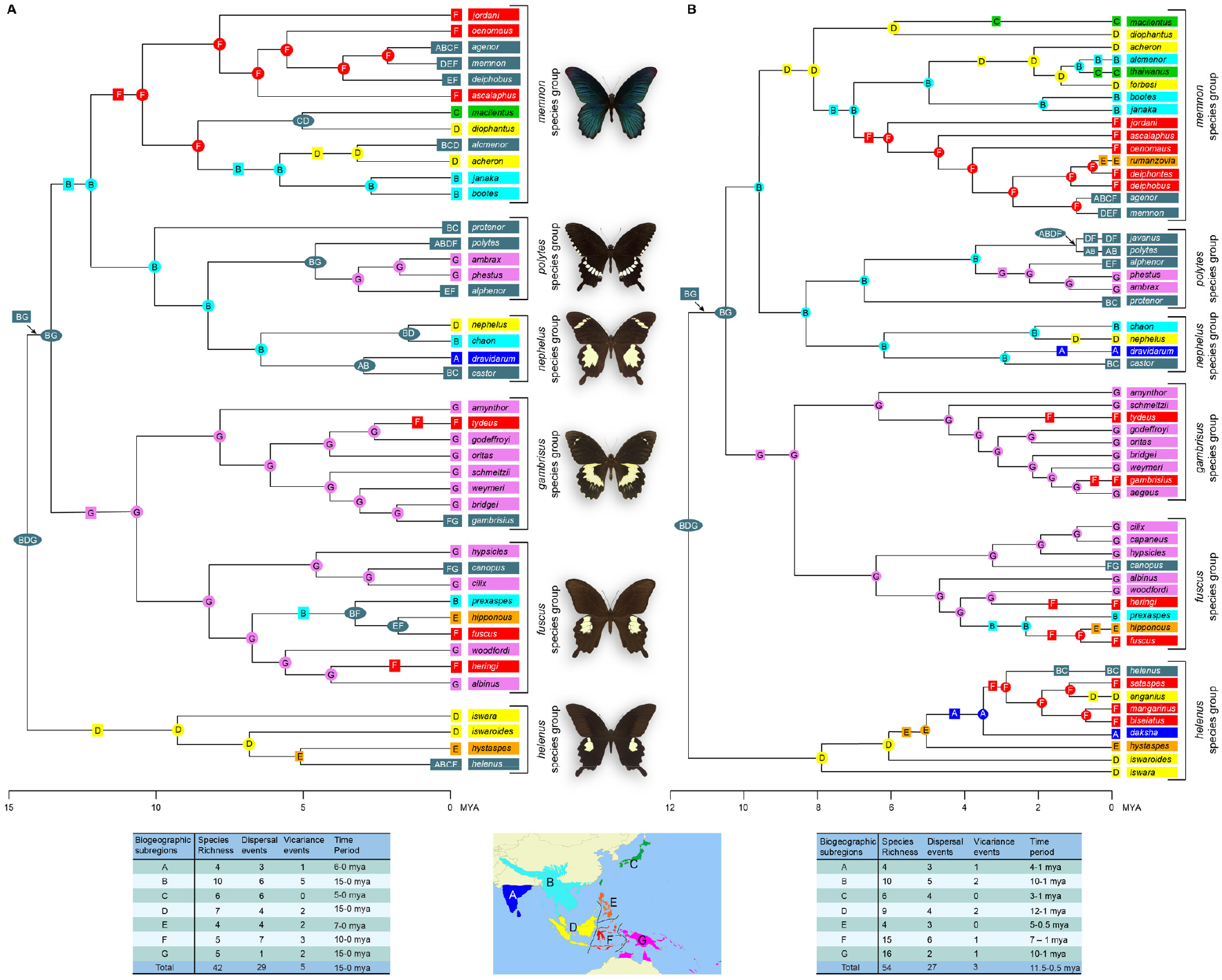
Chronogram showing ancestral area reconstruction using DIVA+J method (the best model, see Table 1) for the Indo-Australian *Menelaides*. **A:** Ancestral area reconstruction for species as identified by species delimitation methods (Fig. 3A). **B:** Ancestral area reconstruction for species as identified by total evidence (Fig. 3B). Letters at internal nodes and their background coloured boxes/circles indicate the reconstructed ancestral area corresponding to respective biogeographic subregions. Letters at the terminal nodes indicate the current geographic distribution of species, with distribution in multiple subregions coloured dark grey. Letters in squares on branches are dispersal or vicariance events, and letters in circles at internal nodes represent estimated ancestral areas. Names of extant species and their distributions are colour-coded and boxed for easy reference. The number of dispersal and vicariance events are summarised in tables at the bottom. A=Peninsular India, B=Continental Asia, C=Japan and Taiwan, D=Sundaland, E=The Philippines, F=Wallacea, and G=Sahul Land.

*Menelaides* was composed of six species groups, in which species assignments to species groups agreed with the traditional classification of Häuser *et al*. (Häuser et al. 2005), except in the case of *P. protenor* (see below) (Fig. 3). However, species hypotheses based on the species delimitation analysis and strongly supported reciprocal monophyly revealed a surprising cryptic species diversity within *Menelaides*, requiring extensive taxonomic revisions (Fig. 3). As per the consensus results of the more stringent species delimitation methods, seven taxa previously treated as subspecies were well supported as distinct genetic clades at the species level, whereas 13 taxa previously treated as species were nested within well-supported species clades, requiring a total of 20 taxonomic changes among the 42 species that were delineated (Fig. 3A, Appendix 1). In comparison, strongly supported reciprocal monophyly of biogeographically cohesive taxa suggested elevation of 14 subspecies to species level, and demotion of seven species to subspecies level, requiring a total of 21 taxonomic changes among the 54 species that were delineated (Fig. 3B, Appendix 1). The cryptic diversity revealed was spread across all the species groups and biogeographic zones (Fig. 3–4), indicating that there was widespread mischaracterisation of species and subspecies in traditional (morphological) taxonomic treatments. Two good cases in point for demotion of species were *P. polymnestor* and *P. inopinatus.* Both these taxa are widely considered to be strongly differentiated species that are endemic to Sri Lanka and peninsular India (*polymnestor*) and the islands of the Maluku province of Indonesia (*inopinatus*). All our analyses showed these to be embedded within *P. agenor* and *P. aegeus*, respectively, with strong support. On the other hand, *P. agenor* and *P. chaon* are widely treated as subspecies of *P. memnon* and *P. nephelus*, respectively, but our phylogenetic analyses (Fig. 3A and 3B) support their elevation to species level. In all, approx. 40% of the delineated species require a taxonomic change by elevating or demoting species and subspecies, which has important implications for the taxonomy and systematics of *Menelaides* (Fig. 3, Appendix 1).

### Molecular Divergence and Species Delimitation in *Menelaides*

Molecular divergence (*p*–distance) between sister clades of species within *Menelaides* varied from <1% to 8% (Fig. 5). However, there was no break in the range of values for molecular divergence between species within *Menelaides*, and subspecies within any species, forming a gradual species-to-subspecies continuum in the range of values of molecular divergence (Fig. 5). Moreover, even the level of molecular divergence of the newly delimited species and/or strongly supported monophyletic clades was well within the range shown by traditionally well-regarded species (red data points in Fig. 5). This indicated that elevation from subspecies to the newly delimited *Menelaides* species was not due to poor performance of the newly developed species delimitation methods. Instead, this suggested that the traditional taxonomic assignments of species and subspecies were not congruent with the level of molecular divergence revealed by our analysis.

### Divergence Time Estimation, Diversification and Biogeography of *Menelaides* in the Indo-Australian Region

In ancestral area reconstruction using DEC, DIVA and BayArea methods, the models with parameter ‘J’ evaluating founder speciation events were chosen in both the likelihood ratio test and the AIC framework (Table 1). The most recent common ancestor of *Menelaides* might have been widespread in the Indo-Australian Region but its precise origin was uncertain due to low probability in the three models: according to the DEC+J (LnL=-135.88) and DIVA+J (LnL=-133.87) models, the ancestor was confined to Sundaland, and according to the BayArea+J model (LnL=-139), it occurred in continental Asia, Sahul Land and Sundaland. According to DIVA+J and DEC+J, there were three vicariance and two dispersal events during the Miocene that led to four distinct *Menelaides* lineages in three distinct biogeographic zones: (1) Sundaland (leading to *helenus* and *memnon* species groups), (2) continental Asia (leading to *memnon*, *polytes* and *nephelus* species groups), and (3) Sahul Land (leading to *gambrisius* and *fuscus* species groups). Thus, these models reconstructed a congruent dispersal/vicariance scenario across the *Menelaides* species tree. Since DIVA+J was chosen as the best model by weighted AICs, we show only DIVA+J reconstruction of ancestral areas and dispersal/vicariance history of *Menelaides* (Fig 4), with remaining models compared in Table 1.

**Table 1:** *Menelaides* ancestral range reconstruction in BioGeoBEARS with three methods: (a) dispersal-extinction-cladogenesis (DEC), (b) dispersal-vicariance analysis (DIVA), and (c) BayArea, along with founder speciation parameter (+J). The ΔAIC compares models with and without J (e.g., DEC vs. DEC+J). The model with the lowest log likelihood score (DIVA + J) is marked bold.

| Models | Log Likelihood | No. of model parameters | AIC | ΔAIC | Dispersal rate (d) | Extinction rate (e) | Founder speciation parameter (J) |
| --- | --- | --- | --- | --- | --- | --- | --- |
| DEC | -156.8 | 2 | 317.6 | 39.8 | 0.0200059 | 1.76E-03 | 0 |
| DEC + J | -135.9 | 3 | 277.8 | 0 | 0.0100188 | 1.00E-12 | 0.07539763 |
| DIVA | -147.1 | 2 | 298.2 | 24.5 | 0.0242666 | 2.00E-09 | 0 |
| **DIVA + J** | **-133.9** | **3** | **273.7** | **0** | **0.0123529** | **4.85E-09** | **0.06099658** |
| BAYAREA | -180 | 2 | 364.1 | 80.1 | 0.0259031 | 2.23E-01 | 0 |
| BAYAREA + J | -139 | 3 | 284 | 0 | 0.0075915 | 1.00E-07 | 0.08948978 |

The diversification of *Menelaides* in the geologically complex Indo-Australian Region was influenced to a great extent by dispersal events, to a lesser extent by vicariance events and range expansions, and this was true across the biogeographic subregions and the six *Menelaides* species groups (Fig. 4). Of the 29 dispersal events, 28 were forward dispersals from ancestral areas to new subregions previously unoccupied by that clade, and only one was a back-dispersal into ancestral areas of that clade. A significant amount of *in-situ* diversification occurred in two subregions: Sahul Land and Wallacea, largely within *memnon*, *gambrisius* and *fuscus* species groups (Fig. 4).

The time of diversification varied considerably between the biogeographic subregions: species diversification occurred much more (10–16 species) near the origin—which broadly corresponds with current centres of diversity—of *Menelaides* over the past 12 million years (inset table in Fig. 4). Diversification occurred much more recently (3–4 million years) and to a much lesser extent (4–6 species) near the periphery of the *Menelaides* distribution, especially in Sri Lanka-Peninsular India and Japan-Taiwan subregions, following more recent dispersals into those subregions.

## DISCUSSION

Considerable advances have taken place in the methods and applications of molecular systematics, phylogenetics, species delimitation methods, and more recently an integration of all of this with phylogenomic data (Hillis et al. 1996; Felsenstein 2004; de Queiroz 2007; McCormack et al. 2013; Misof et al. 2014; Rannala 2015; Kawahara et al. 2019; Hime et al. 2021). Much progress has been made since the importance of universal molecular markers and the use of mito-nuclear evidence, phylogenetic methods and integrative taxonomy was emphasized (Caterino et al. 2000; Wahlberg and Wheat 2008; Dupuis et al. 2012; Papakostas et al. 2016; Yang and Rannala 2017). As a result, taxonomy and the age-old problems of defining species and subspecies are beginning to be tackled with these new methods and tools. The use of molecular systematics and integrative taxonomy hold considerable promise for solving taxonomic problems such as those posed by *Papilio* and other diverse taxa, but efforts are just catching up to do so (Kunte et al. 2011, 2019; Andersen et al. 2014; Dupuis and Sperling 2015, 2016; Toussaint et al. 2015; Matos‐Maraví et al. 2019). This may help taxonomists define species in manners that are systematically robust, and that will also provide well-informed species hypotheses for evolutionary biologists to study biogeography, population divergence, potential evolution of reproductive barriers, and diversification.

On this background, *Papilio* is an illuminating example. Systematics of *Papilio* in light of the significant polytypy and even more striking morphological (wing colour pattern) differentiation has traditionally been a problem that divided taxonomic opinion and practice. The phylogenetic and molecular systematic results presented above are among the strongest applications of modern methods to resolve age-old taxonomic issues of polytypy and species definitions in *Papilio*. The strong branch supports from a mito-nuclear dataset used in our phylogeny also suggest that our molecular sampling is likely adequate for this work. The use of larger genome-wide molecular datasets would be an important improvement on the existing work, especially where our work shows a disagreement between species delimitation methods (Fig. 3A) and phylogenetically strongly supported, biogeographically meaningful monophyly (Fig. 3B). Specifically, it will be important to attempt to resolve the following four cases with phylogenomic data. First, in our phylogenetic analysis, *P. protenor* was placed at the base of the *polytes* species group, although it is usually believed to be a member of the *memnon* species group on morphological grounds. Second, our analysis placed the island taxon *merapu* inside the predominantly mainland monophyletic group that we referred to as *P. agenor*, instead of inside the predominantly island clade (*P. memnon*) to which it might belong. Third, *P. alphenor* was placed in the subgroup (*alphenor* (*phestus*, *ambrax*)) when it is expected to be in the *polytes* + *javanus* subgroup based on wing patterns and mimetic female polymorphism. Finally, the cryptic species revealed by well-supported monophyly and widely allopatric distributions in the *helenus* species group are striking. Note that these phylogenetically well-supported findings are based on dense taxon sampling and inclusion of multiple samples of each taxon traditionally treated as species (SI Table 1), and a molecular dataset that hardly had any gaps, minimizing the possibility that these phylogenetic placements were artefacts. Phylogenomic approaches may be able to further resolve these unexpected results, in additional to providing further support for diversification in the highly diverse *memnon* species group.

The complex polytypy and phylogenetic distinctiveness described above is expected to unfold over an equally complex geological land/seascape. The Indo-Australian Region forms such a geologically complex landscape, setting stage for among the most complicated biogeographic scenarios for biotic diversification in the world (Lohman et al. 2011; Bacon et al. 2013; Brown et al. 2013). Its tall, ragged mountains, deep oceanic fissures and strong oceanic currents have influenced—and in some places prevented—dispersal and divergence of distinctive lineages within this region, giving rise to a unique biogeographic mix of various mainland and island faunas (Wallace 1860, 1876; Simpson 1961; Mayr 1965; Lohman et al. 2011; Bacon et al. 2013; Brown et al. 2013). The various biogeographic barriers responsible for this isolation in some areas and mix in others have geologically formed from plate tectonics and geo-climatic cycles. These geo-climatic cycles, along with the history of dispersals and vicariance in a fragmented landscape, bear strong signatures on the patterns of diversification of *Papilio* swallowtails, which we summarize below. The synthetic view of the evolutionary history of Cenozoic period of the Indo-Australian Region is still emerging, when studies like ours with detailed phylogenies and fine-scale biogeographic analyses across the region will be crucial for understanding diversification processes in the landscape.

Sea level changes associated with geo-climatic cycles, and movement of the continental plates, have been critical in the formation of the Indo-Australian Region and the evolution of biodiversity therein. By early Miocene (25 mya), continental plates, many island arcs and major biogeographic barriers such as the Makassar Strait and the Wallace’s Line had assumed their present positions (Hall 1998). Mid-Miocene period is also recognised as the ‘climatic optimum’ in this area as the warming phase occurred 17–15 mya and led to the extensive growth of tropical forests throughout the Sundaland. Subsequently, there was cooling that continued until 6 mya and led to the contraction of tropical forests. This was again followed by a warming phase that continued until ~3.2 mya. During this period, tectonic activity and uplift also affected both the positioning of several islands (e.g. the Philippines) and the shape and extent of individual islands (Hall 2002). These warm-wet climate cycles and geological changes in the Miocene-Pliocene appear to have played an important role in shaping some of the evolutionary divergences in the Indo-Australian Region across taxa including plants, birds, butterflies and mammals (Meijaard 2004; Lohman et al. 2011; Brown et al. 2013).

*Menelaides* species time tree with ancestral area reconstruction suggested that *Menelaides* started diversifying in the mid-to late-Miocene (approx. 11 mya; Fig. 4) with a widespread ancestor possibly across Sundaland, continental Asia and Sahul shelf. At the time, Sundaland was still one landmass (largely Borneo) and connected to the Asian mainland, while Sumatra probably consisted of an island arc and Java was still mostly submerged (Meijaard 2004). The initial three vicariance and two dispersal events led *Menelaides* diversification across the Indo-Australian Region. Four major *Menelaides* clades leading to six species groups had started to diversify in three distinct biogeographic zones (continental Asia, Sundaland and Sahul Land) towards the late Miocene (10–8 mya; Fig. 4).

After these initial vicariance events, dispersals across biogeographic subregions dominated *Menelaides* diversification, coupled to a lesser extent with *in situ* speciation in these biogeographic subregions (inset tables in Fig. 4). On the whole, *Menelaides* in the Sahul Land largely diversified through *in situ* speciation (9–2mya), although there was inter-island dispersal within this subregion. Diversification in the Sundaland, Wallacea and continental Asia appears to have been shaped by a combination of dispersals and *in situ* speciation from mid-Miocene to Pliocene (11–2.6 mya). This finding supports the emerging view that diversification in Wallacea and Asia was shaped by events in the Miocene and Pliocene (Stelbrink et al. 2012). Two isolated oceanic island chains (the Philippines and Japan) received *Menelaides* relatively recently (mostly 5–0.5 mya) through repeated dispersals from adjoining biogeographic subregions. Similarly, in peninsular India and Sri Lanka—which were far-flung from the centre of *Menelaides* diversity and therefore acted as habitat islands on the western edge of the Oriental Region—*Menelaides* dispersed recently (5–1 mya), and then diversified through some *in situ* speciation. Throughout this process of dispersal and *in situ* speciation across all the biogeographic subregions and in all the species groups, a predominant and peculiar pattern emerges: no two sister species of *Menelaides* occur on the same islands or within the same subregions without dispersal barriers between them. Indeed, very few species even within a species group occur within the same fine-scale island groups, and most of them occur in allopatry across major dispersal barriers, often separated on mainland and islands, or across major island groups. Thus, the predominant pattern of diversification and current species ranges is that *Menelaides* diversified almost exclusively in allopatry.

Detailed phylogenetic analyses revealed that species diversity in *Menelaides* was mischaracterised by traditional taxonomic treatments of this iconic butterfly subgenus. The unexpectedly high level of cryptic clade diversity, which was spatially strongly structured in the mainland-island mosaic, showed widespread reciprocal monophyly of biogeographically meaningful clades, in almost all species groups. This work provides critical insights into the diversification process of this important group of butterflies and also a useful case study of the biogeography, morphological diversification and speciation in this dense cluster of endangered biodiversity hotspots of the Indo-Australian Region.

## AUTHOR CONTRIBUTIONS

KK conceived and supervised the study; JJ designed and performed phylogenetic and biogeographic analyses; JJ and KK wrote the manuscript.

## ACKNOWLEDGEMENTS

We thank Adam Cotton and Felix Sperling for generous contributions of specimens/tissue samples and advice on *Papilio* and molecular systematics; Manas Samant, Ravi Umadi, Rhucha Vatturkar, Anupama Prakash and the NCBS Sequencing Facility for DNA sequencing; and Felix Sperling, Adam Cotton and Rohit Naniwadekar for comments on the manuscript. New Indian material used in this work was collected under research and voucher specimen collection permits issued by the state forest departments in Kerala (permit no. WL 10-3781/2012 dated 18/12/2012, and GO (RT) No. 376/2012/F&WLD dated 26/07/2012) and Nagaland (permit no. CWL/GEN/240/522-39 dated 14 August 2012), which is now deposited in the Biodiversity Lab Research Collections at NCBS. Additional specimens were provided by Laurie Wills.

## Appendix

**Appendix 1:**
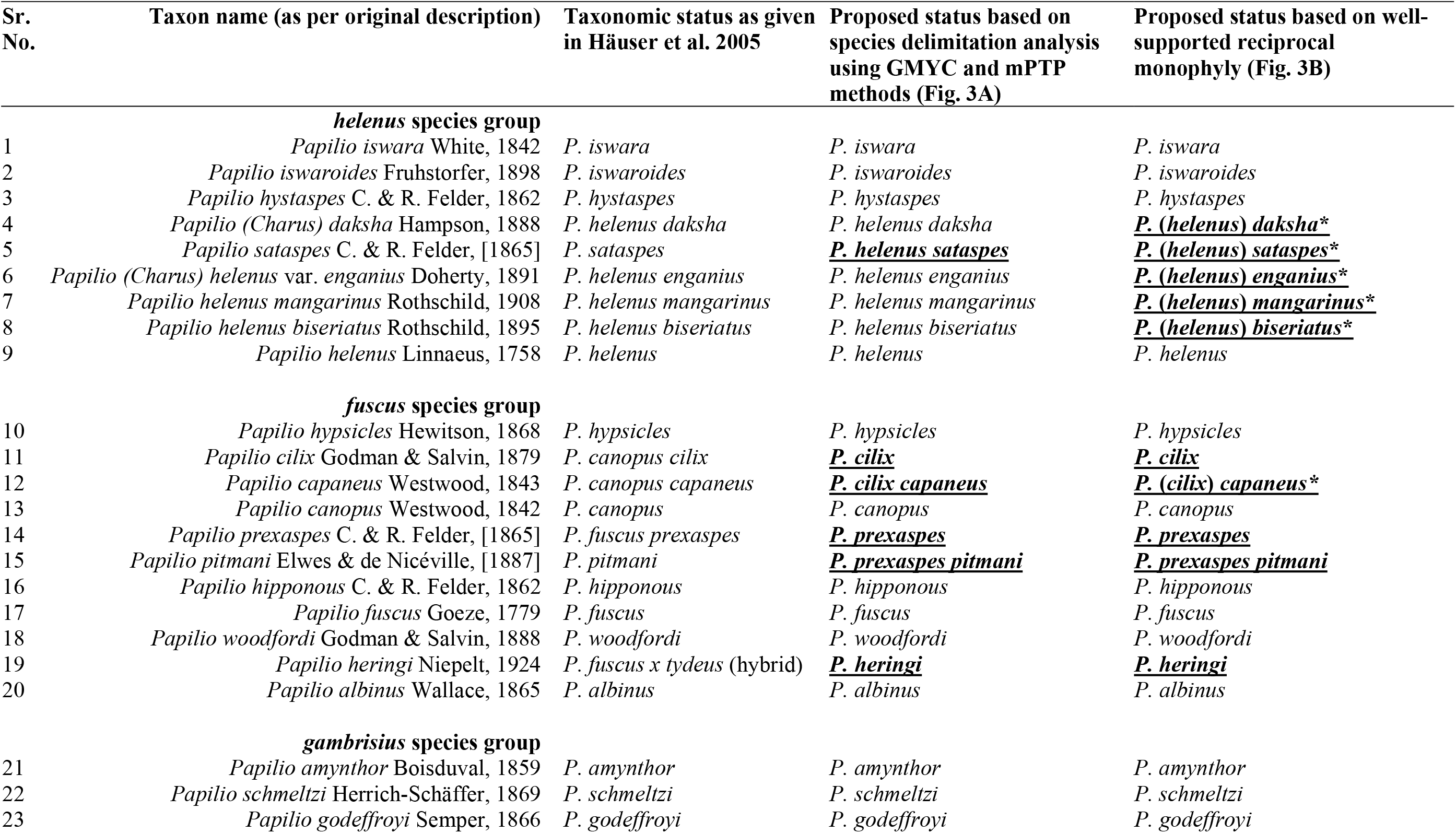
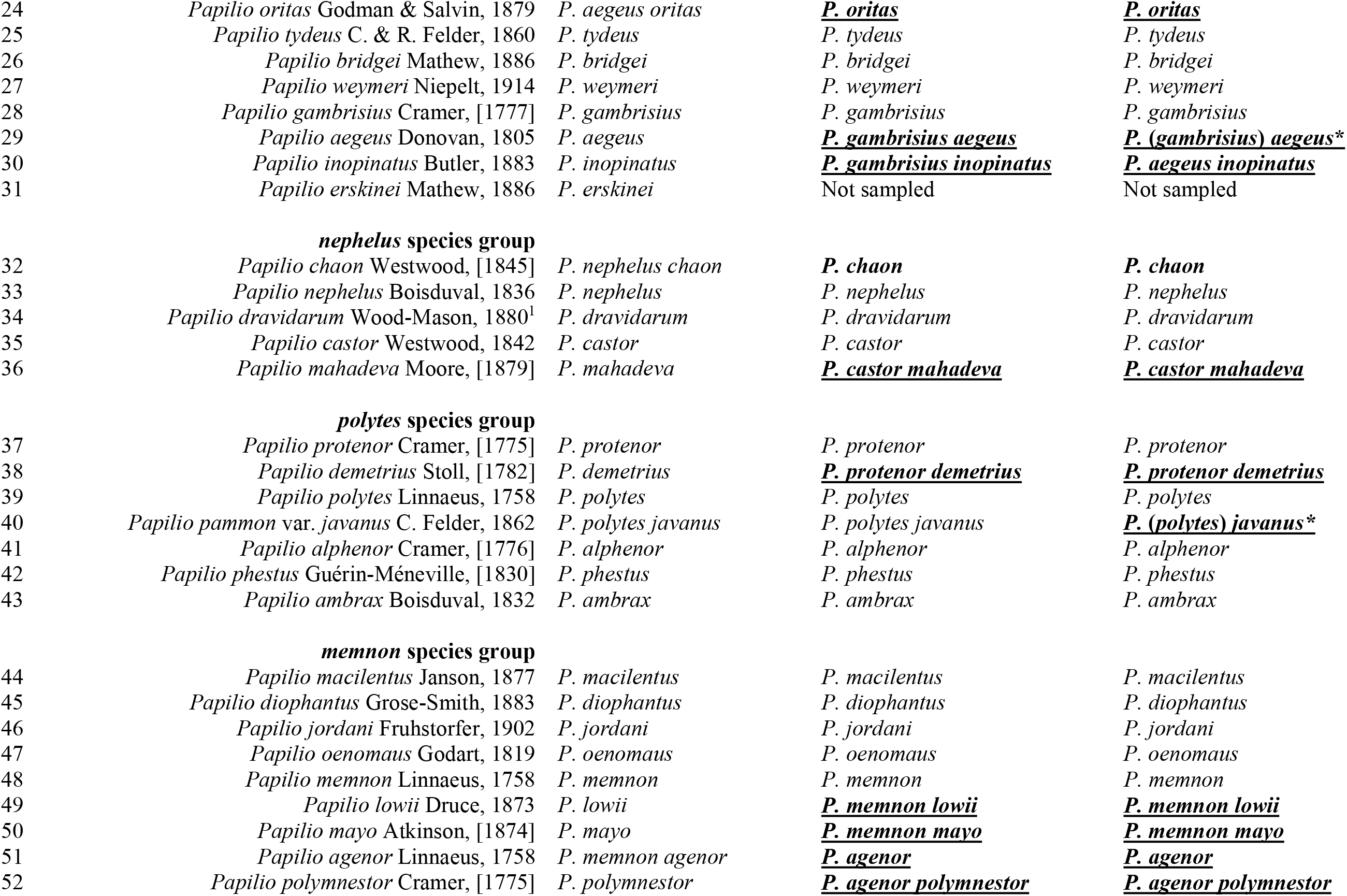
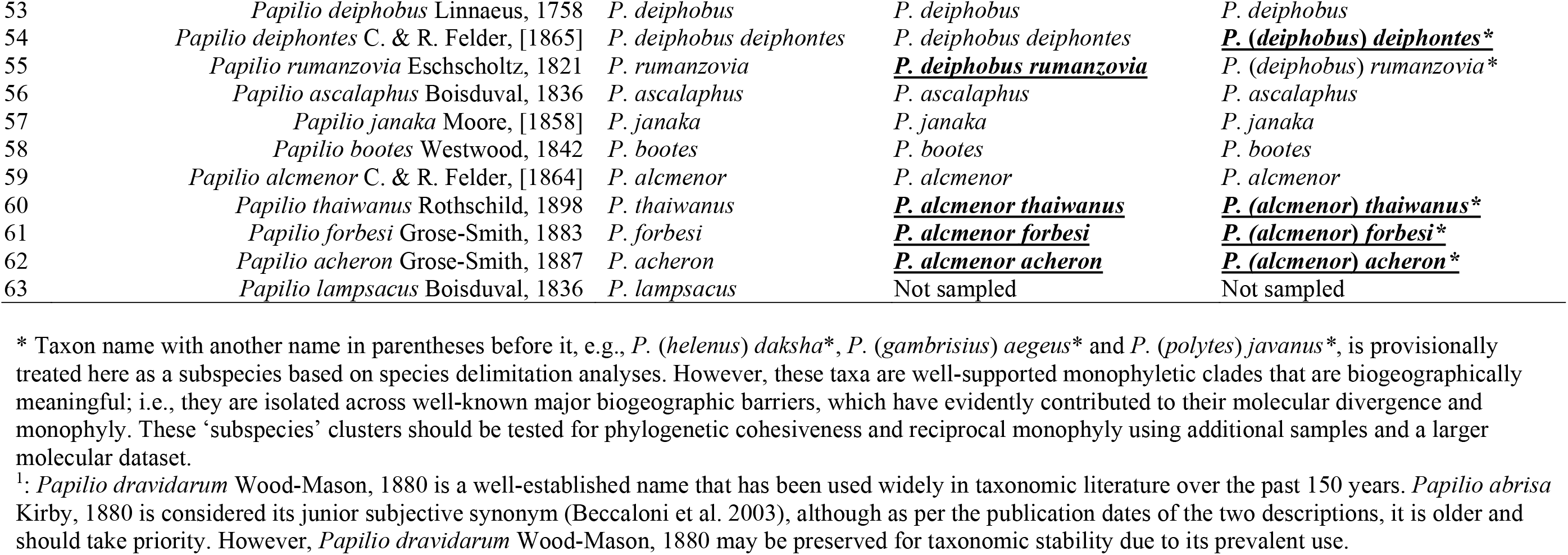
Species checklist of *Menelaides* based on the analyses presented here. Taxon names that are underlined and marked bold are taxonomic changes with respect to the latest taxonomic checklist of *Menelaides* (Häuser et al. 2005).

## SUPPLEMENTARY TABLES AND FIGURES

**SI Table 1:** This table follows classification of *Papilio* species as inferred from our molecular phylogenetic analyses. In other words, these are current hypotheses about species and higher taxonomic classification of *Papilio* (also see Appendix 1). Given under gene names are Genbank accession numbers. See Excel sheet, SITable1_TaxaDetails.xlsx.

**SI Table 2:** Information about the primers and loci used in this study. See Word document, SITable2_PrimersInfo.docx.

**SI Table 3:** Phylogenetic support (posterior probability) from the species delimitation methods— Generalized Mixed Yule Coalescent (GMYC) and Multi-rate Poisson Tree Process model (mPTP)—to delineate *Menelaides* species. Species names follow Appendix 1. See Excel sheet, SITable3_SpeciesDelimitationResults.xlsx.

**SI Figure 1:**
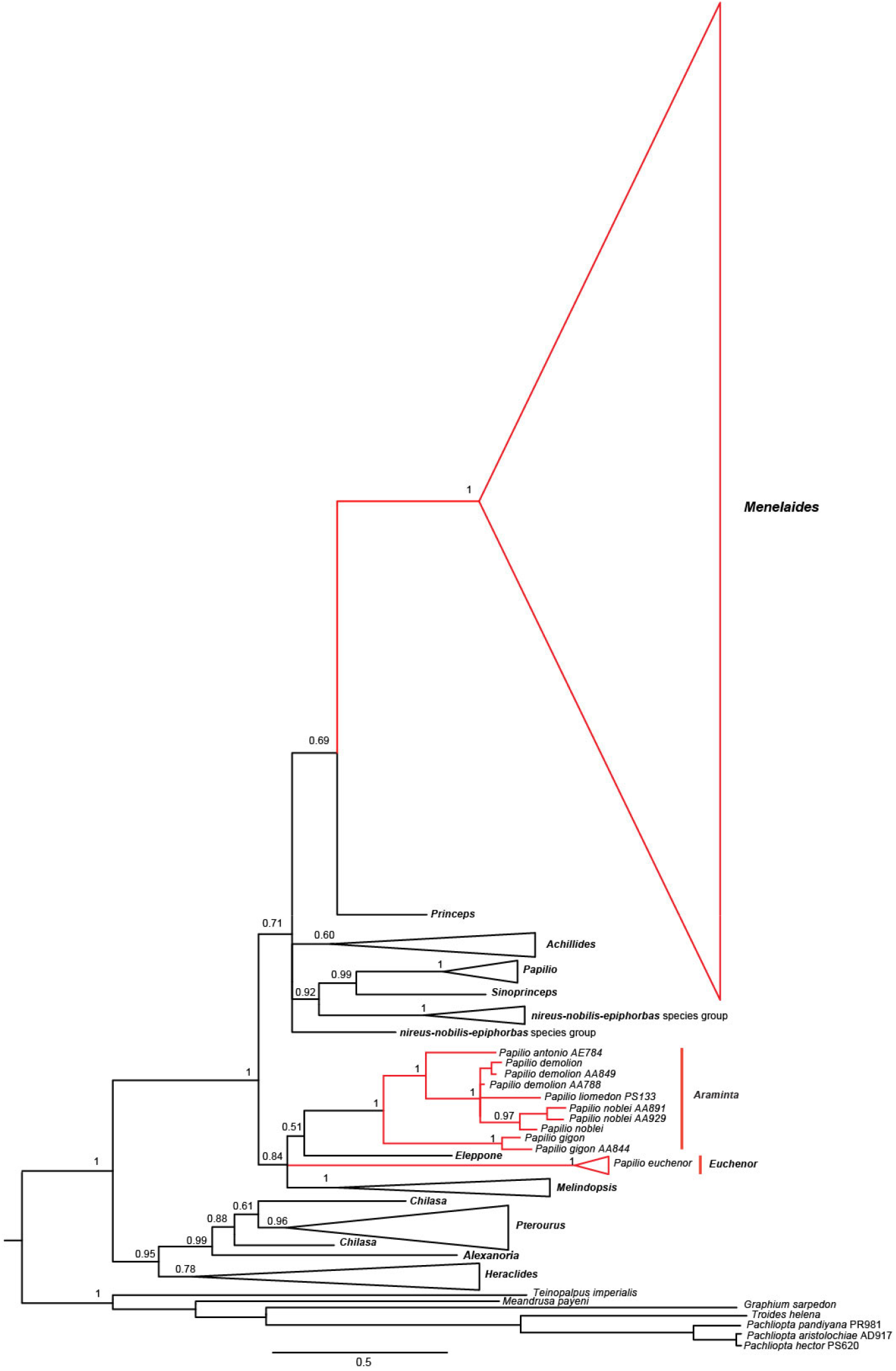
A partitioned Bayesian phylogram of *Papilio* based on two nuclear markers (*EF1-alpha* and *wingless*). Bayesian posterior probability is indicated at each node. Taxonomic position and monophyly of *Menelaides* as fixed by our phylogenetic analyses is marked red. Taxa that were previously treated under *Menelaides* but are now found to be outside the subgenus are also marked red.

**SI Figure 2:**
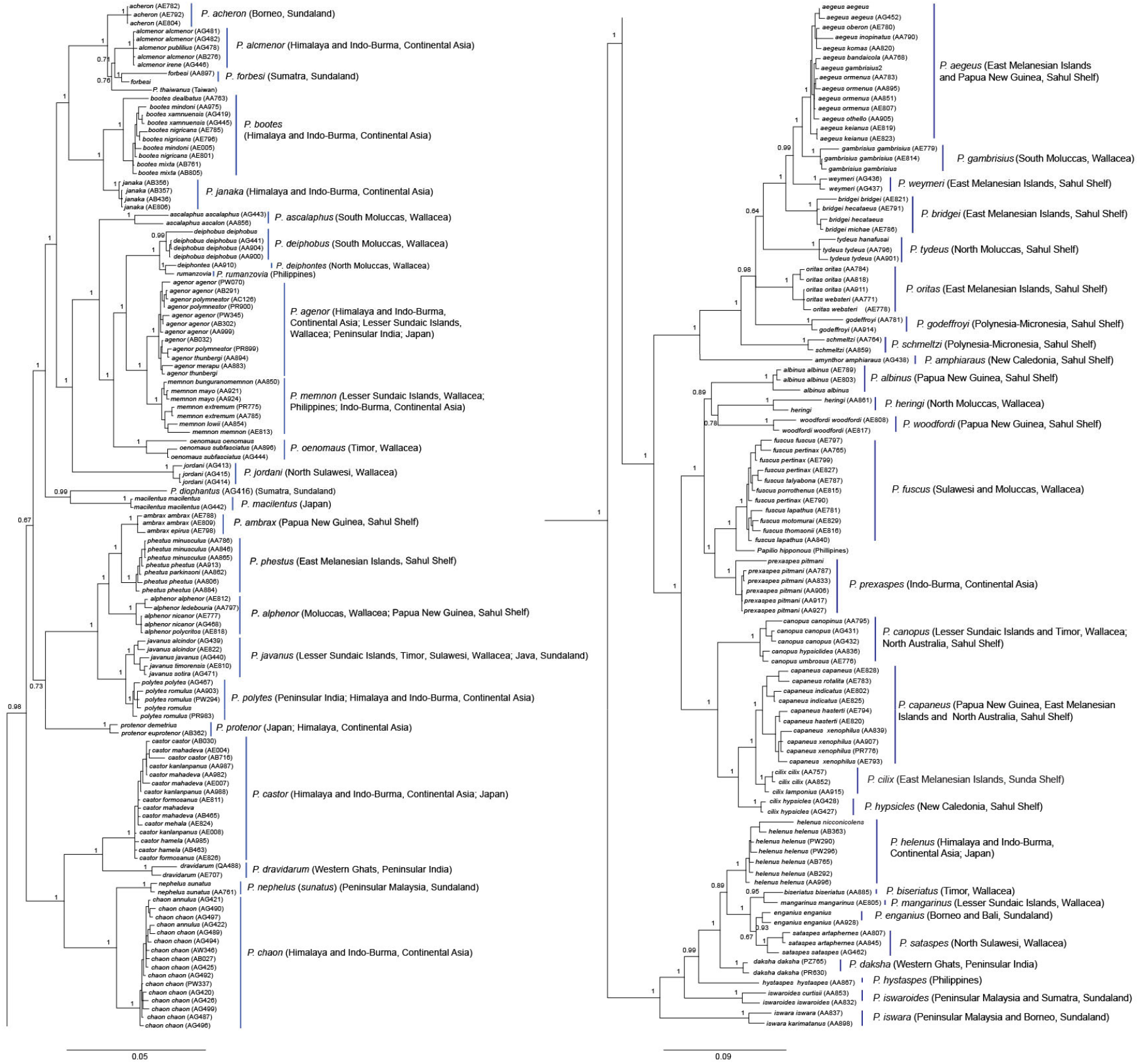
A detailed Bayesian phylogram of *Menelaides* species of the Indo-Australian Region based on three mitochondrial and two nuclear markers along with voucher codes, taxa names and geographic distribution.

